## Supplemental Material for "Concerted modification of nucleotides at functional centers of the ribosome revealed by single-molecule RNA modification profiling"

### Single-molecule modification profiling of ribosomal RNA reveals concerted modification at functional locations in the ribosome

Andrew D. Bailey IV, Jason Talkish, Hongxu Ding, Haller Igel, Alejandra Durán, Shreya Mantripragada, Benedict Paten, and Manuel Ares, Jr.

#### Supplemental Materials

##### Supplemental Figures

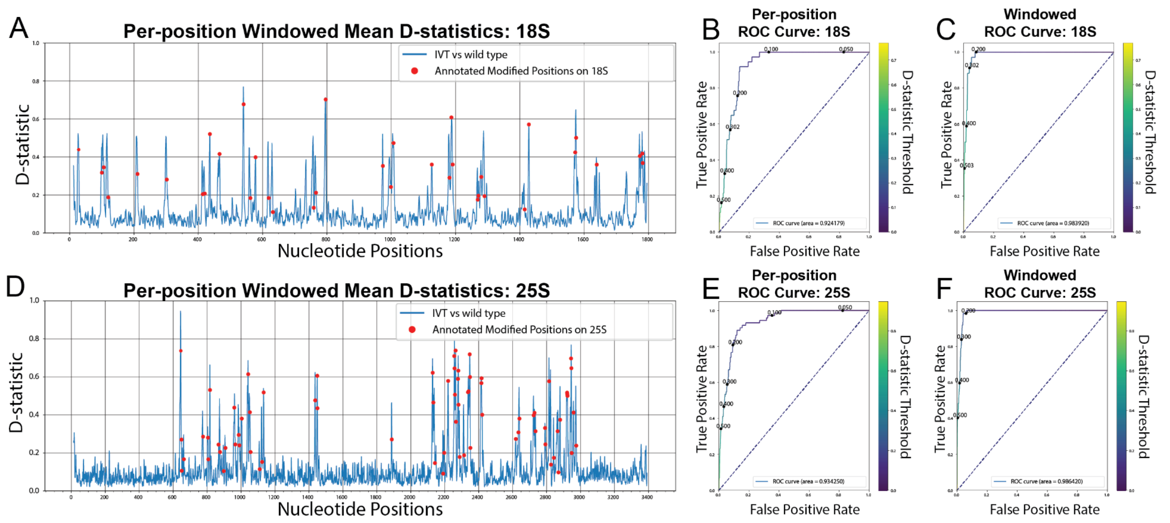


**Figure 1-figure supplement 1: De-novo detection of modifications using Tombo.** (A,D) Per position, window averaged D-statistic plots from Tombo's sample compare method for yeast 18S (A) and 25S (D) rRNA (Stoiber et al. 2016). The blue line represents the difference between the per-position distributions of the IVT sample vs the wild type sample. The red markers are the location of each annotated modification on the corresponding rRNA (Taoka et al. 2016). (B,E) Per-position modification calling AUROC (Area Under the Receiver Operating Characteristic) curves for Tombo's sample compare method comparing the IVT sample vs a wild type sample for yeast 18S (B) and 25S (E) rRNA. (C,F) Modification window calling AUROC curves for Tombo's sample compare method comparing the IVT sample vs the wild type sample for yeast 18S (C) and 25S (F) rRNA (see Methods).

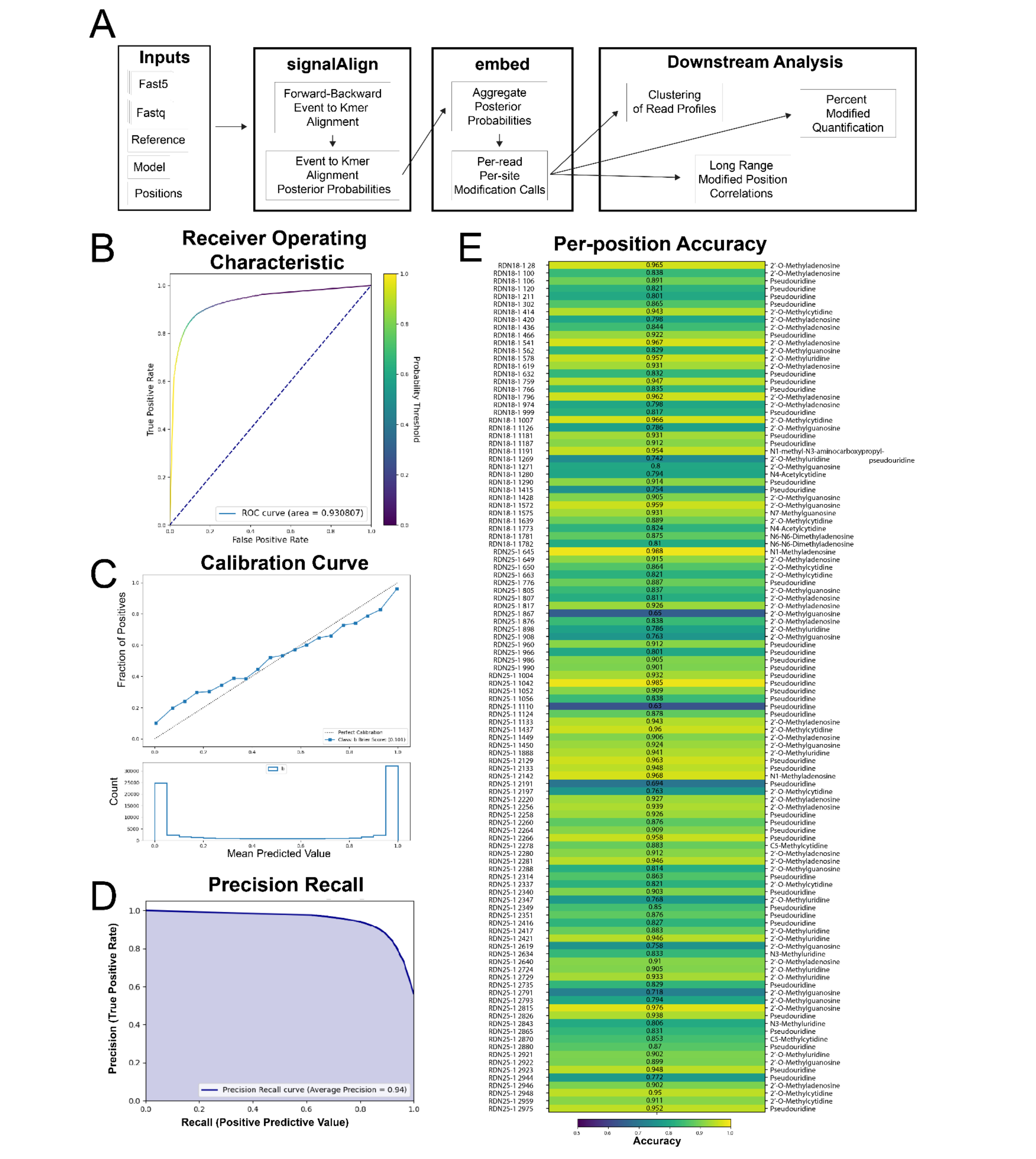


**Figure 1-figure supplement 2: SignalAlign pipeline overview, overall accuracy metrics from testing data and per-position model accuracy.** (A) Analysis pipeline. (B-E) Testing accuracy metrics of the final model of supervised training. Both training protocol and testing metrics are described in detail in Methods. (B) Receiver operating characteristic (ROC) curve and area under the ROC (0.93). (C) Calibration curve showing the fraction of true positives for several ranges of probabilities. The brier score (0.101) is a metric for determining how well a model is calibrated. (D) Precision-recall curve. (E) Per-position accuracy with corresponding modification annotation for each position (Taoka et al. 2016).

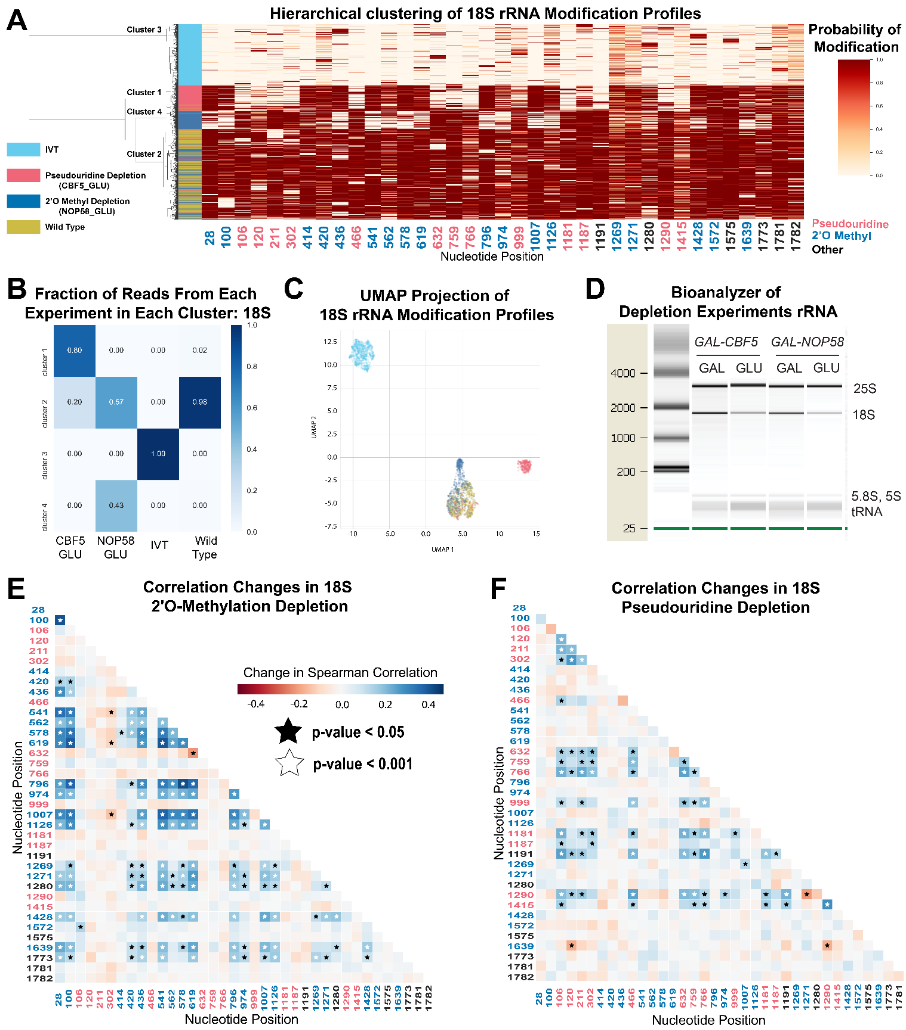


**Figure 1-figure supplement 3**: **Clustering and correlation analysis of depletion experiment modification profiles in 18S rRNA.** (A) Hierarchical clustering of 18S yeast rRNA modification profiles of IVT, wild type, and both pseudouridine and 2'O methyl depletion experiments. (B) Fraction reads from IVT, wild type and both depletion experiments (CBF5_GAL, NOP58_GAL) in each cluster of 18S rRNA. (C) UMAP visualization of 18S yeast rRNA modification profiles of IVT, wild type, and both pseudouridine and 2'O methyl depletion experiments. UMAP color scheme is the same as the labels in panel A. (D) Bioanalyzer of comparing levels of 18S and 25S rRNA in galactose-grown samples (CBF5_GAL, NOP58_GAL) compared to glucose-grown samples (CBF5_GLU, NOP58_GLU). (E/F) Change in Spearman correlations of 25S reads in 2'O methyl depletion (E) and pseudouridine depletion (F) when compared to wild type. Stars represent significant changes when compared to wild type correlation and significantly different from zero correlation (see Methods).

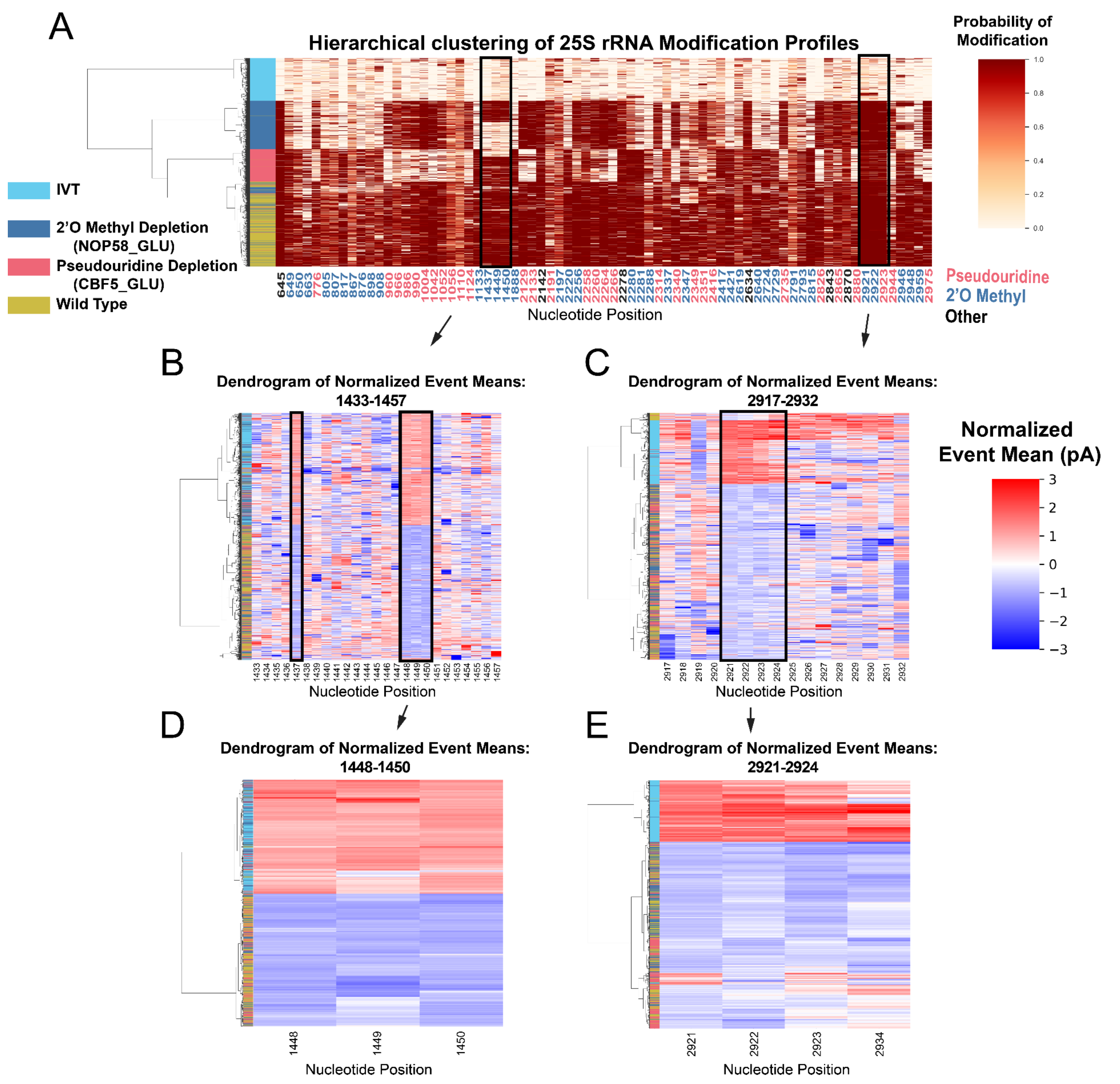


**Figure 1-figure supplement 4: Clustering of underlying events to search for patterns of modification in the pseudouridine and 2'O methyl depletion experiments.** (A) Hierarchical clustering of 25S yeast rRNA modification profiles of IVT, wild type, and both pseudouridine and 2'O methyl depletion experiments (1000 reads in each experiment). Each row represents a full length single read, each column represents a modified nucleotide and the scale represents the probability of being modified. (B-E) Hierarchical clustering of normalized event means aligned to the reference sequence from IVT, wild type, and both pseudouridine and 2'O methyl depletion experiments covering positions 1433 to 1457 (B), 2917-2932 (C), 1448-1450 (D), and 2921-2924 (E) (see Methods and Supplemental Note 1).

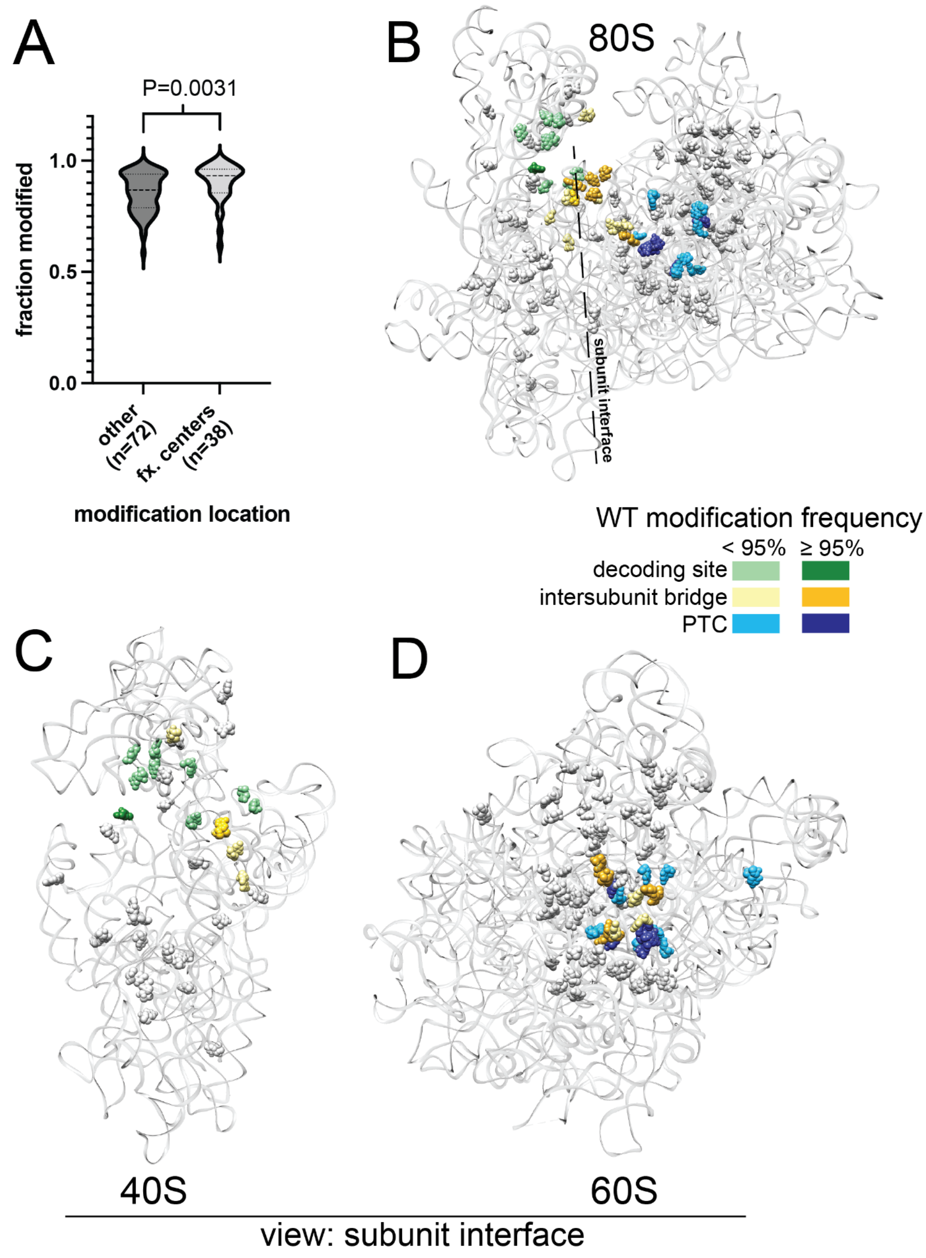


**Figure 1-figure supplement 5: Analysis of yeast rRNA modification frequency in relation to functional centers of the ribosome.** (A) Distribution of fraction modified for positions within or not within the functional centers of yeast rRNA. Distribution means are significantly (p-value=0.0031) different via a two-sided Mann-Whitney U-test. (B-D) Crystal structure model of wild type *S. cerevisiae* 80S (B), 40S (C) and 60S (D) rRNA highlighting modification frequency within functional centers. PDB: 4V88 (Ben-Shem et al. 2011).

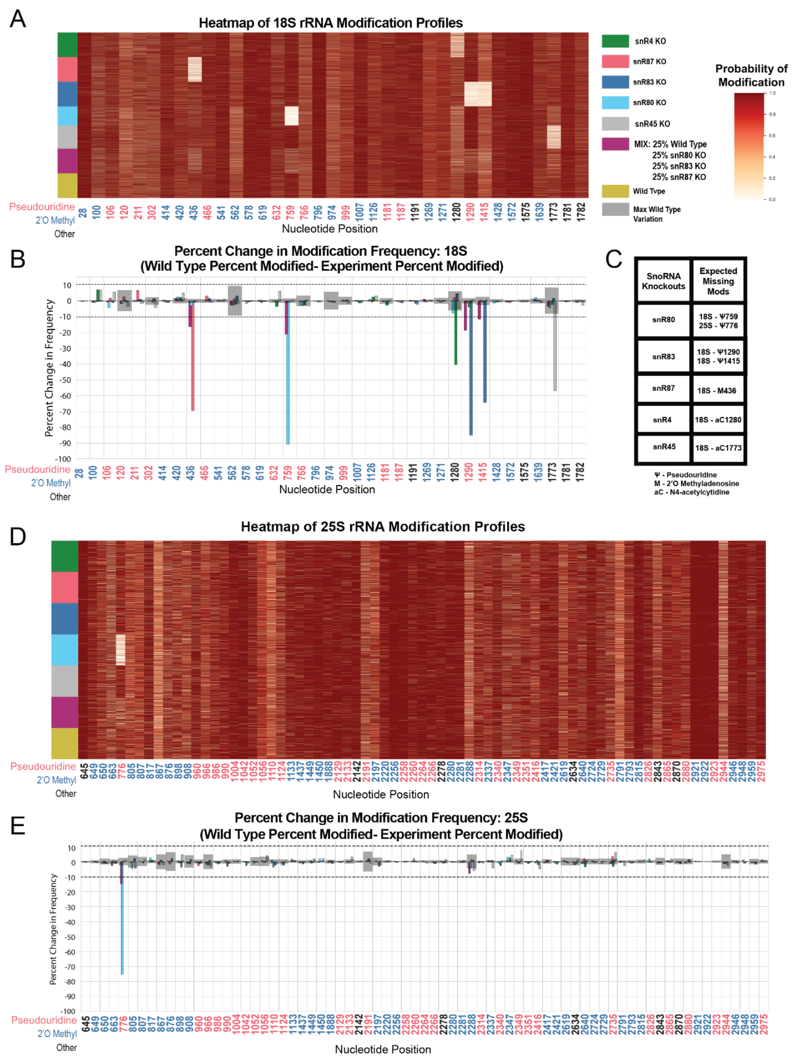


**Figure 2-figure supplement 1: Heatmaps and percent modification change of snoRNA knockout and mixture experiments.** (A) Heatmap of wild type, mixed sample, snR80Δ, snR83Δ, and snR87Δ, snR45Δ and snR4Δ modification profiles of 18S (1000 reads in each experiment). Each row represents a full length single read, each column represents a modified nucleotide and the scale represents the probability of being modified. (B) Mixed sample, snR80Δ, snR83Δ, and snR87Δ, snR45Δ and snR4Δ 18S percent change in modification frequency when compared to wild type. Gray bars indicate the variance of wild type rRNA modification at each position and the black dotted lines represent the maximum variance found at any position. (C) Table of snoRNAs knocked down with the corresponding expected knocked down modifications. (D) Heatmap of wild type, mixed sample, snR80Δ, snR83Δ, and snR87Δ, snR45Δ and snR4Δ modification profiles of 25S (1000 reads in each experiment). (E) Mixed sample, snR80Δ, snR83Δ, and snR87Δ, snR45Δ and snR4Δ 25S percent change in modification frequency when compared to wild type.

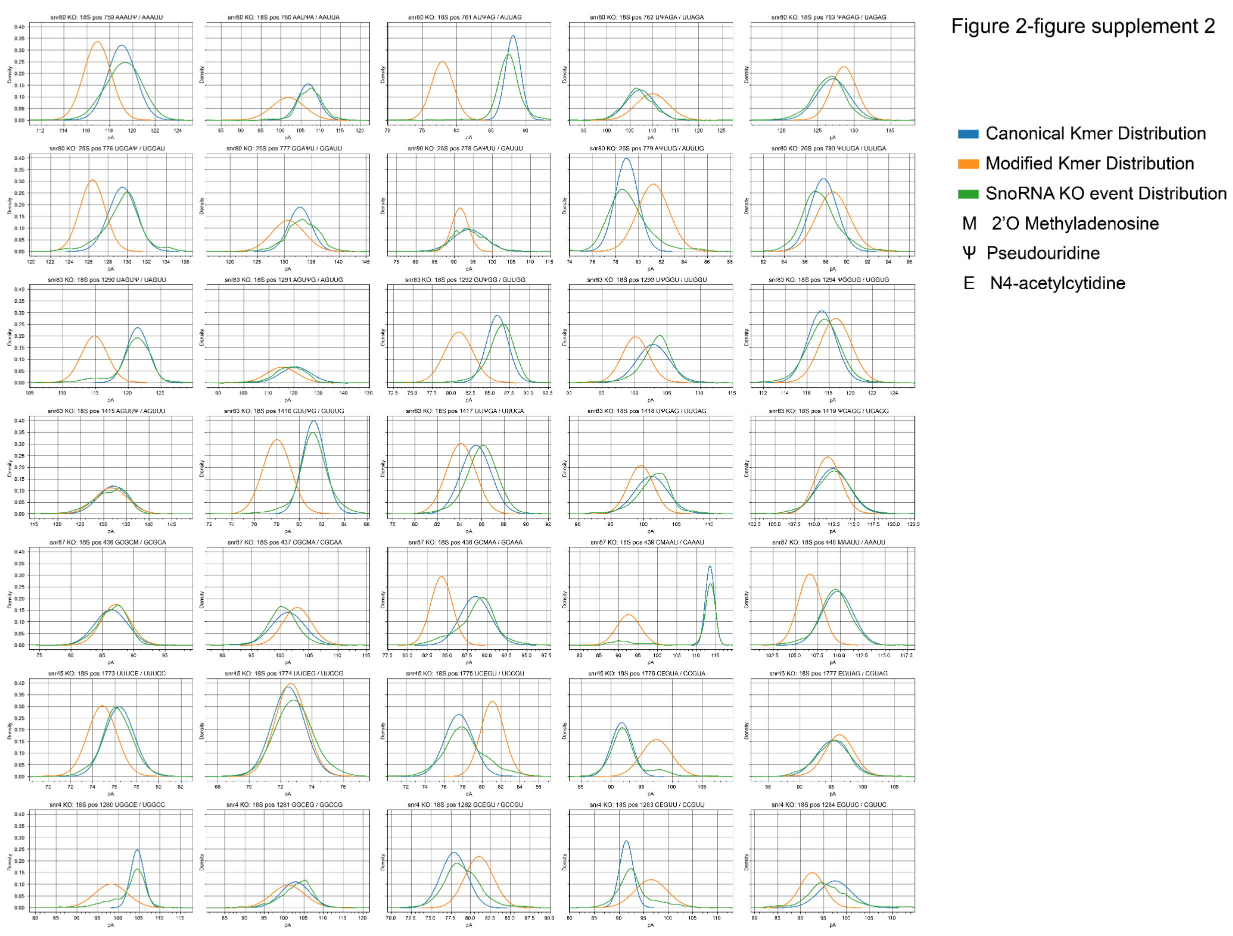


**Figure 2-figure supplement 2: Kmer distribution comparison between snoRNA knockout kmer distributions and the trained model kmer distributions.** Each figure has the model's canonical kmer distribution, the model's modified kmer distribution and the corresponding snoRNA knockout kernel density estimate (KDE) of all events aligned to that position (see Methods). The rows show kmers covering position 759 in 18S from snR80Δ, position 776 in 25S from snR80Δ, position 1290 in 18S from snR83Δ, position 1415 in 18S from snR83Δ, position 436 in 18S from snR87Δ, position 436 in 18S from snR87Δ, position 1773 in 18S from snR45Δ and position 1280 in 18S from snR4Δ.

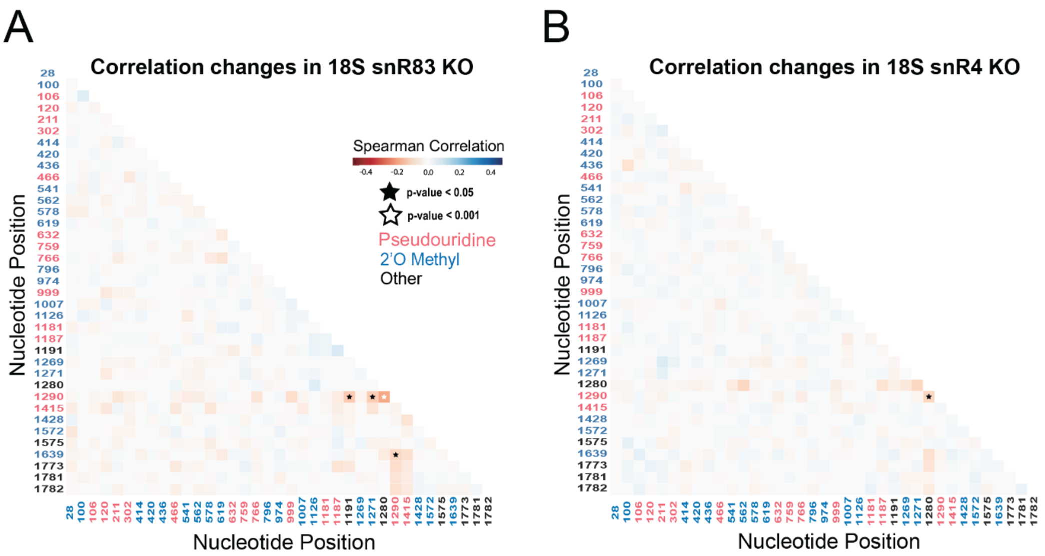


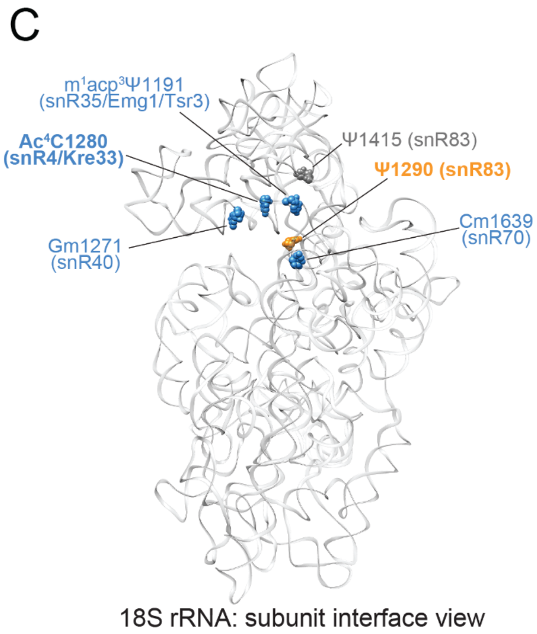


D

**snR83 KO (Ψ1290)** | **95** | **9**

| correlated position | snoRNA/ enzyme | %mod WT | %mod mut. |
|---|---|---|---|
| m¹acp³Ψ1191 | snR35/ Emg1/Tsr3 | 93 | 91 |
| Gm1271 | snR40 | 88 | 90 |
| ac⁴C1280 | snR4/ Kre33 | 76 | 83 |
| Cm1639 | snR70 | 86 | 86 |

**snR4 KO (ac⁴C1280)** | **76** | **32**

| correlated position | snoRNA/ enzyme | %mod WT | %mod mut. |
|---|---|---|---|
| Ψ1290 | snR83 | 95 | 90 |

**Figure 2-figure supplement 3: Clustering and correlation analysis of snoRNA KO experiment modification profiles in yeast 18S rRNA.** (A/B) Change in Spearman correlations of 18S reads in snR83 KO (A) and snR4 KO (B) when compared to wild type. Stars represent significant changes when compared to wild type correlation and significantly different from zero correlation (see Methods). (C) Crystal structure model of wild type *S. cerevisiae* 18S rRNA highlighting significant, concerted positions. PDB: 4V88 (Ben-Shem et al. 2011). (D) Table of percent modification of sites of interest in wild type and snoRNA KO experiments.

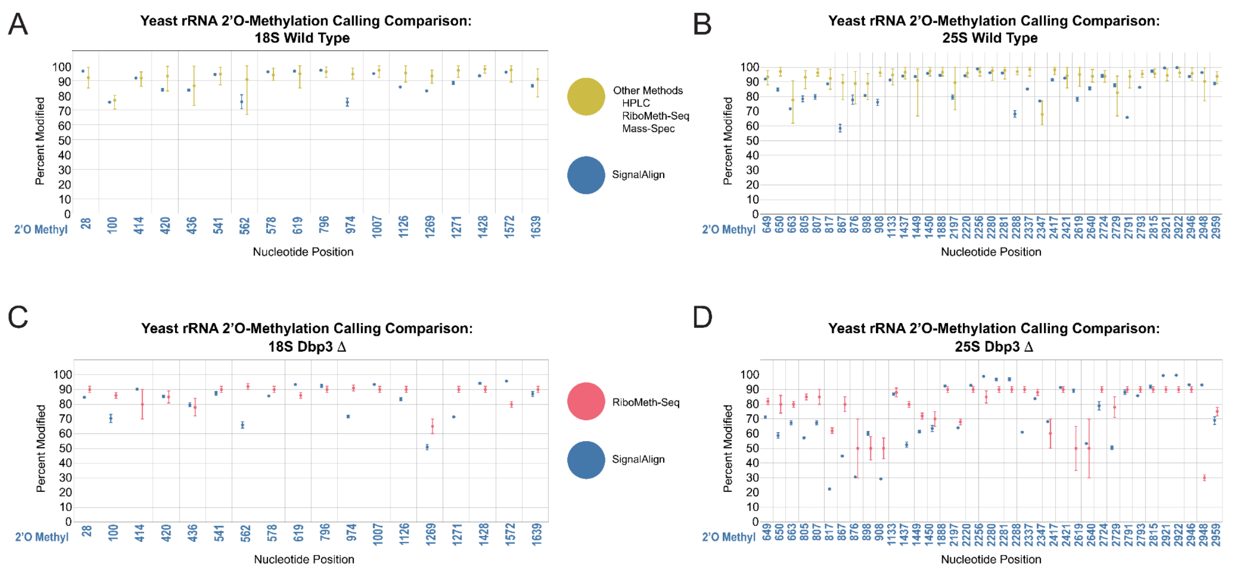


**Figure 2-figure supplement 4: Comparison of rRNA 2'O-methylation calling from other modification detection techniques and signalAlign modification detection.** (A-B) Comparison between the range of modification percentages called via mass spectrometry (Taoka et al. 2016), HPLC (Yang et al. 2016), and two RiboMeth-seq approaches (Birkedal et al. 2014; Marchand et al. 2016) vs signalAlign modification percentages of wild type yeast in 18S (A) and 25S (B). (C-D) Comparison between RiboMeth-seq modification percentages (Aquino et al. 2021) and signalAlign modification percentages for the Dbp3 knockout strain in 18S (C) and 25S (D) yeast rRNA. For the combination of several detection approaches, we calculated the minimum, maximum and mean modification percentage from the four papers. For all plots, error bars represent the minimum or maximum percent modification called and circles represent the mean modification percentage.

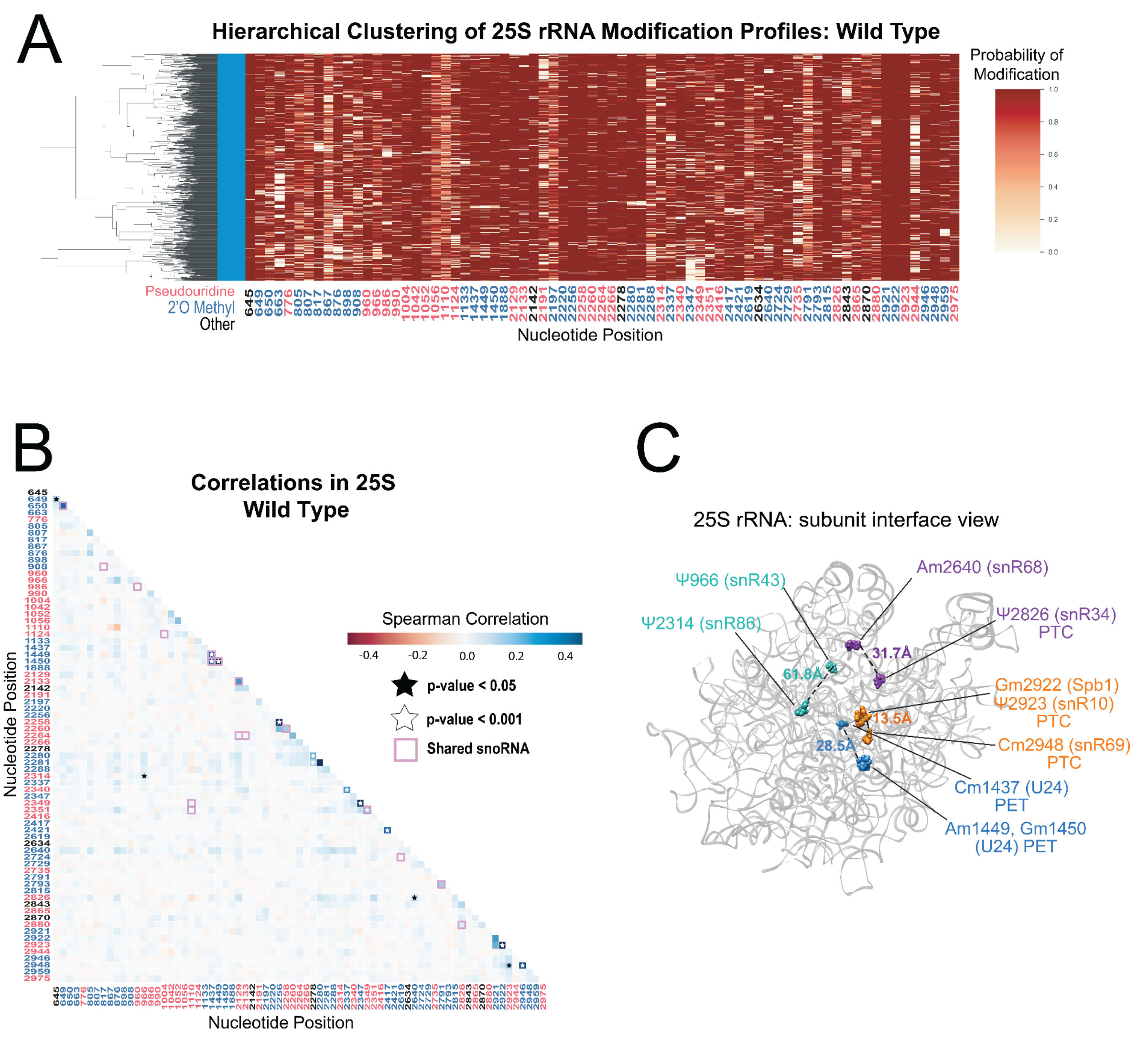


**Figure 2-figure supplement 5: Yeast 25S rRNA modification profile clustering and correlation analysis.** (A) Hierarchical clustering of 25S yeast rRNA modification profiles from wild type yeast (1000 reads). Each row represents a full length single read, each column represents a modified nucleotide and the scale represents the probability of being modified. (B) Wild type Spearman correlation of 25S wild type reads. Stars represent significantly different concerted positions compared to IVT and significantly different from zero correlation. (C) Crystal structure model of wild type *S. cerevisiae* 25S rRNA highlighting significant, concerted positions. PDB: 4V88 (Ben-Shem et al. 2011).

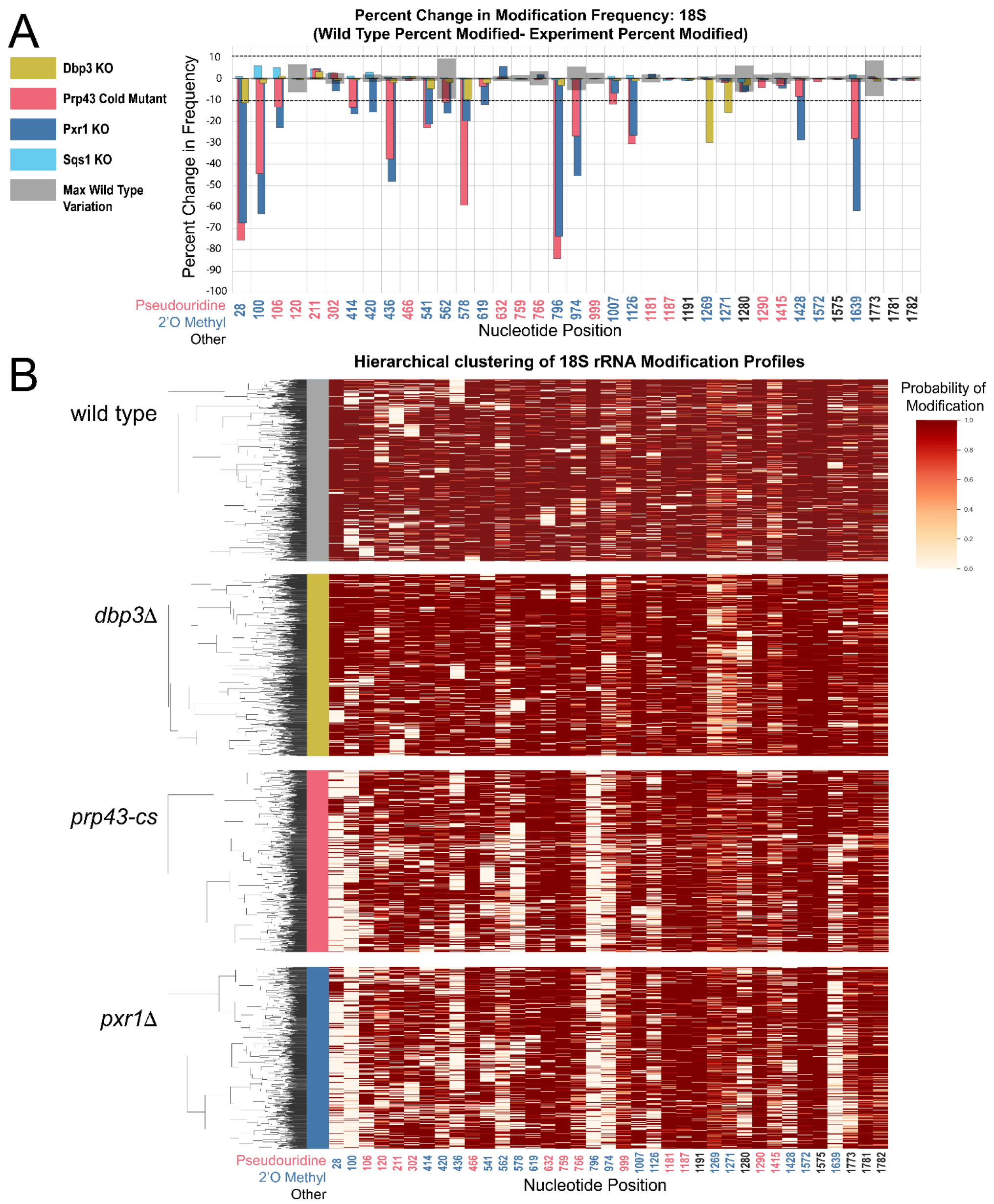


**Figure 3-figure supplement 1: Clustering of 18S rRNA modification profiles and percent change in modification frequency upon mutation of helicases Dbp3 and Prp43 and G-patch proteins Pxr1 and Sqs1.** (A) Barplots of the difference between wild type modification frequency and *dbp3*Δ, *prp43-cs*, *pxr1*Δ, and *sqs1*Δ modification frequencies in 18S yeast rRNA. Gray bars indicate the variance of wild type rRNA modification at each position and the black dotted lines represent the maximum variance. (B) Hierarchical clustering of 18S yeast rRNA modification profiles from wild type, *dbp3*Δ, *prp43-cs*, and *pxr1*Δ (1000 reads in each experiment). Each row represents a full length single read, each column represents a modified nucleotide and the scale represents the probability of being modified.

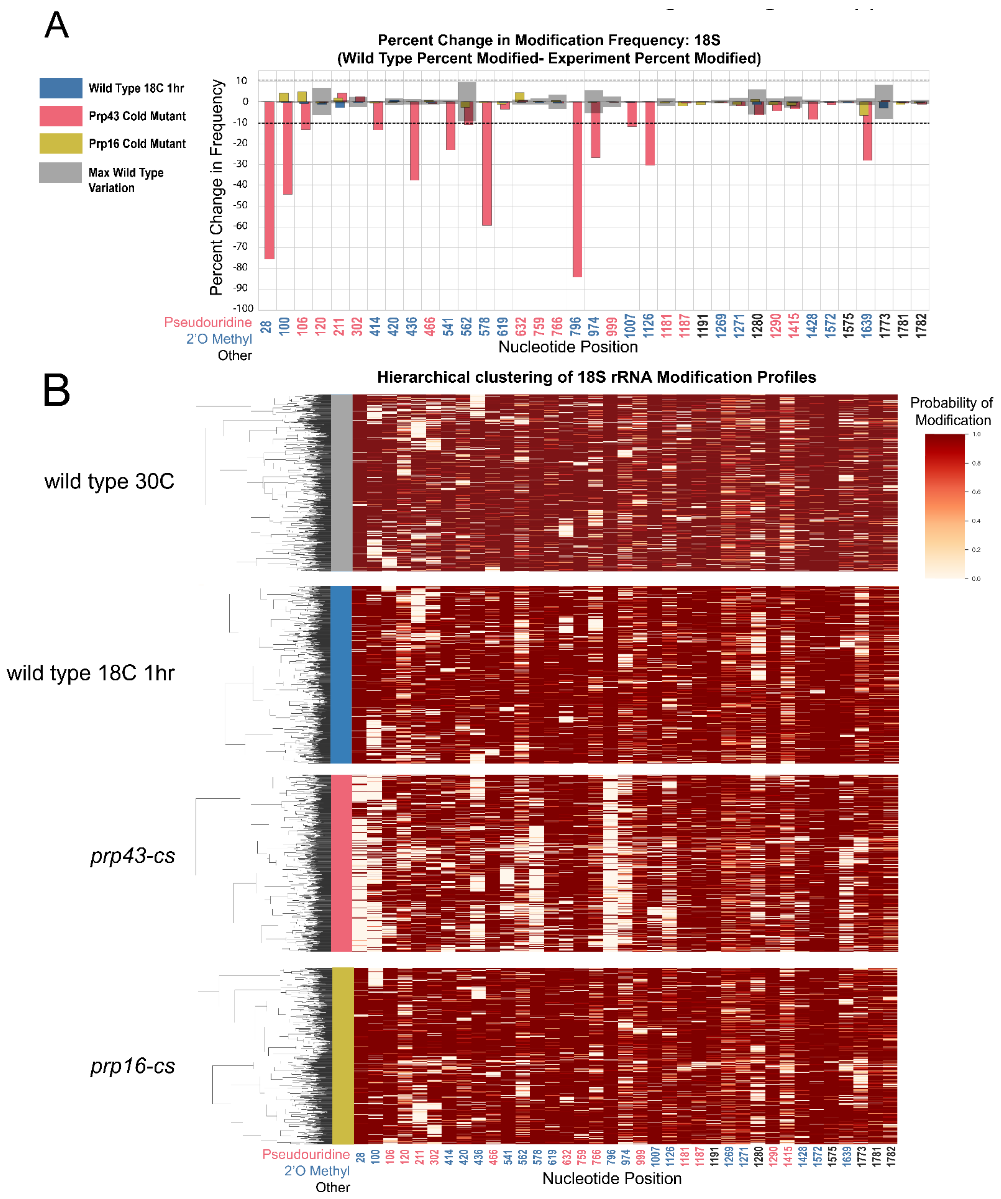


**Figure 3-figure supplement 2: Clustering of 18S rRNA modification profiles and percent change in modification frequency upon mutation of helicases Prp43 and Prp16, compared to wild type controls grown at 30 °C or shifted to 18 °C for 1 hour.** (A) Barplots of the difference between 18S yeast rRNA modification frequency of wild type cells grown at 30 °C and wild type, *prp43-cs*, and *prp16-cs* cells shifted to 18 °C for 1 hour. Gray bars indicate the variance of wild type rRNA modification at each position and the black dotted lines represent the maximum variance. (B) Hierarchical clustering of 18S yeast rRNA modification profiles from wild type 30 °C, wild type 18 °C, *prp43-cs*, and *prp16-cs* (1000 reads in each experiment). Each row represents a full length single read, each column represents a modified nucleotide and the scale represents the probability of being modified.

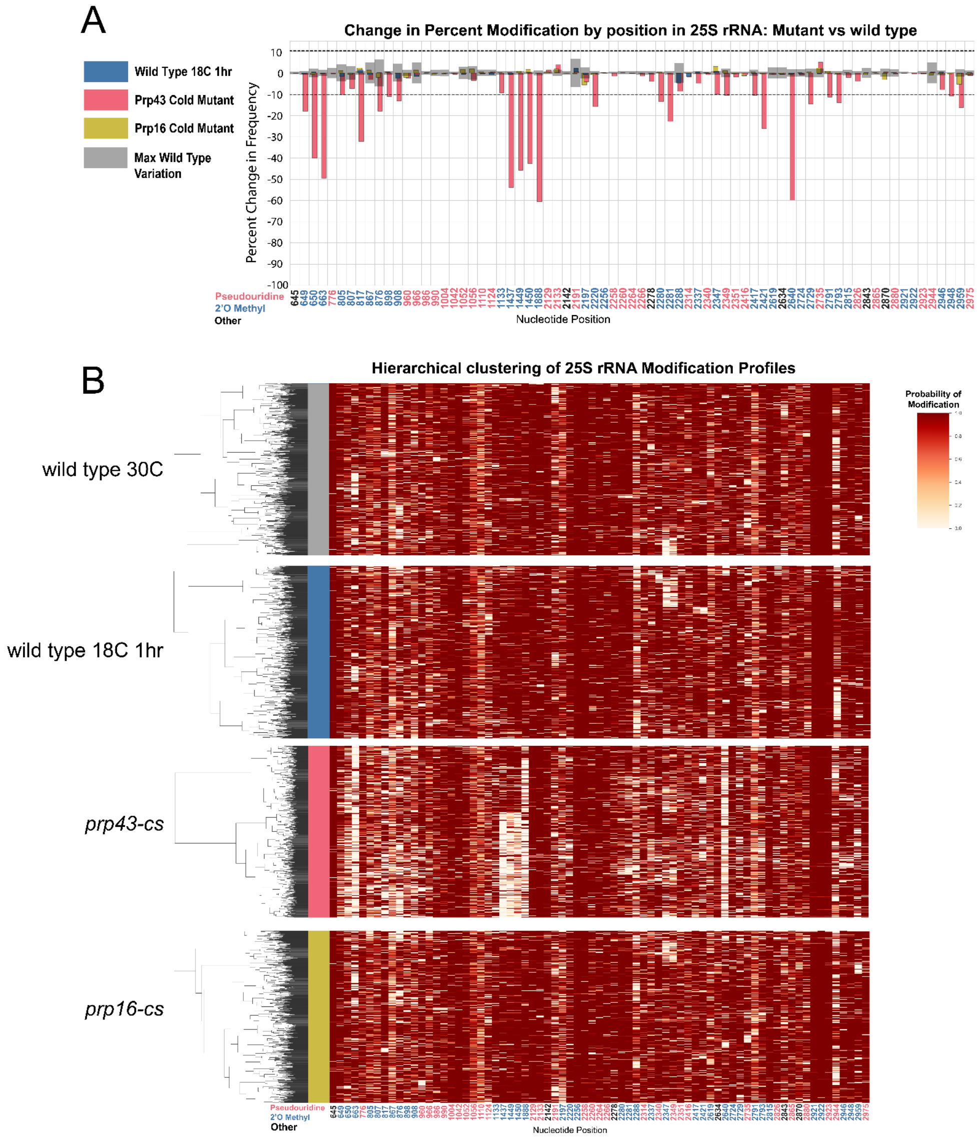


**Figure 3-figure supplement 3: Clustering of 25S rRNA modification profiles and percent change in modification frequency upon mutation of helicases Prp43 and Prp16, compared to wild type controls grown at 30 °C or shifted to 18 °C for 1 hour.** (A) Barplots of the difference between 25S yeast rRNA modification frequency of wild type cells grown at 30 °C and wild type, *prp43-cs*, and *prp16-cs* cells shifted to 18 °C for 1 hour. Gray bars indicate the variance of wild type rRNA modification at each position and the black dotted lines represent the maximum variance. (B) Hierarchical clustering of 18S yeast rRNA modification profiles from wild type 30 °C, wild type 18 °C, *prp43-cs*, and *prp16-cs* (1000 reads in each experiment). Each row represents a full length single read, each column represents a modified nucleotide and the scale represents the probability of being modified.

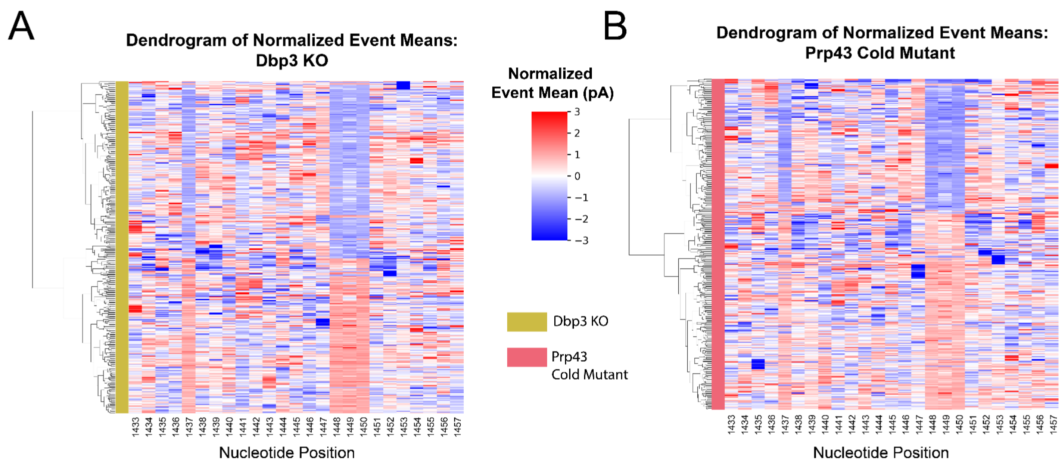


**Figure 3-figure supplement 4: Clustering of underlying events to search for patterns of modification in the Dbp3 KO and Prp43 cold mutant.** Hierarchical clustering of aligned standardized events from Dbp3 KO (A) and Prp43 cold mutant (B) covering the events from positions 1431 to 1455 (see Methods). These positions cover the 3 2'O ribose methylations guided by the snoRNA U24 at positions 1437, 1449 and 1450.

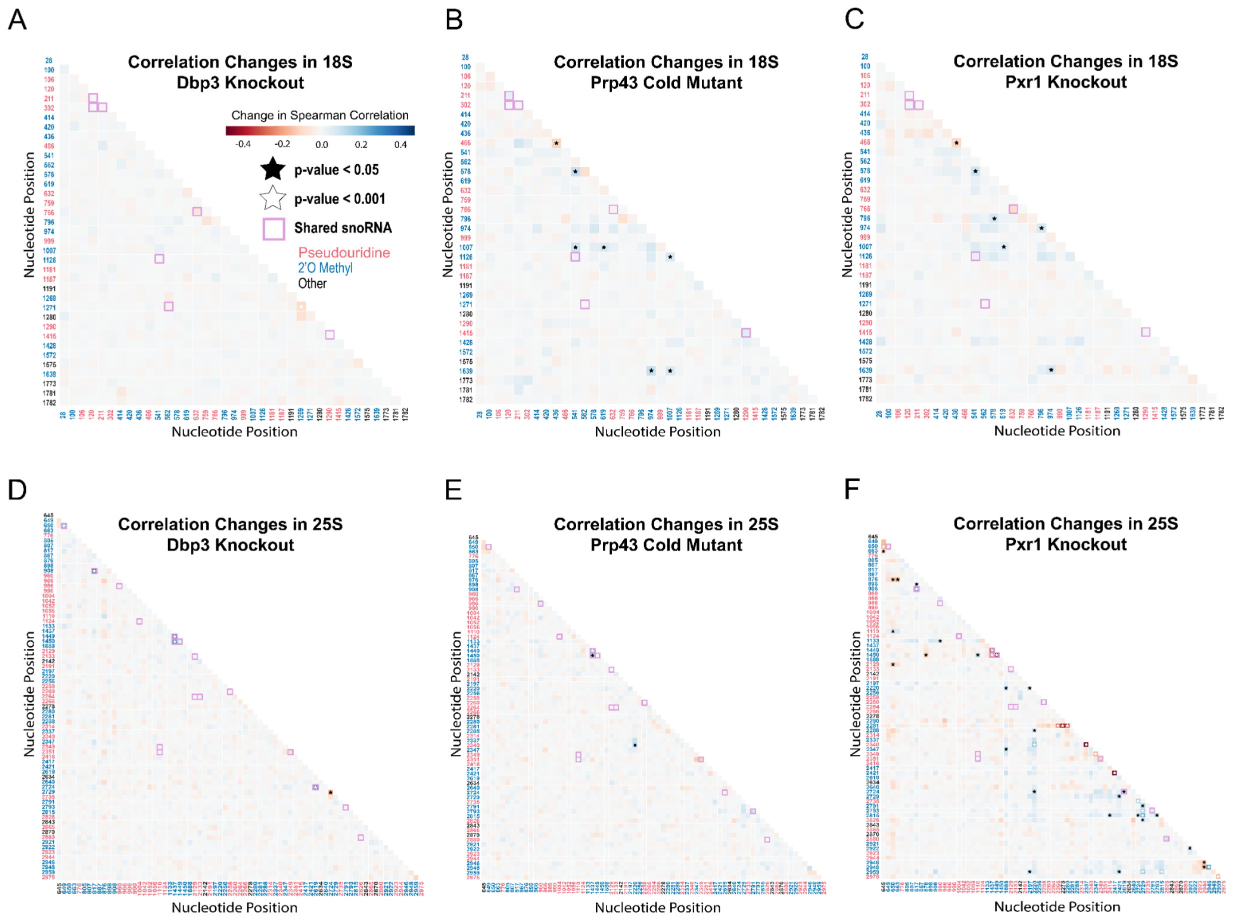


**Figure 4-figure supplement 1: Correlation analysis of *dbp3*Δ, *prp43-cs*, and *pxr1*Δ.** Change in Spearman correlations of 18S (A-C) and 25S (D-E) reads in *dbp3*Δ (A/D), *prp43-cs* (B/E), and *pxr1*Δ (C/F) when compared to wild type. Stars represent significant changes when compared to wild type correlation and significantly different from zero correlation.

Ding, Hongxu, Andrew D. Bailey, Miten Jain, Hugh Olsen, and Benedict Paten. 2020. “Gaussian Mixture Model-Based Unsupervised Nucleotide Modification Number

Detection Using Nanopore-Sequencing Readouts." Edited by Jonathan Wren. *Bioinformatics* , no. June (June): 1–7.

Lapeyre, Bruno, and Suresh K. Purushothaman. 2004. "Spb1p-Directed Formation of Gm2922 in the Ribosome Catalytic Center Occurs at a Late Processing Stage." *Molecular Cell* 16 (4): 663–69.

Marchand, Virginie, Florence Blanloeil-Oillo, Mark Helm, and Yuri Motorin. 2016. "Illumina-Based RiboMethSeq Approach for Mapping of 2′-O-Me Residues in RNA." *Nucleic Acids Research* 44 (16): e135–e135.

Marcus H Stoiber, Joshua Quick, Rob Egan, Ji Eun Lee, Susan E Celniker, Robert Neely, Nicholas Loman, Len Pennacchio, and James B Brown. 2016. "De Novo Identification of DNA Modifications Enabled by Genome-Guided Nanopore Signal Processing." *biorXiv*. https://doi.org/10.1101/094672.

Taoka, Masato, Yuko Nobe, Yuka Yamaki, Yoshio Yamauchi, Hideaki Ishikawa, Nobuhiro Takahashi, Hiroshi Nakayama, and Toshiaki Isobe. 2016. "The Complete Chemical Structure of Saccharomyces Cerevisiae rRNA: Partial Pseudouridylation of U2345 in 25S rRNA by snoRNA snR9." *Nucleic Acids Research* 44 (18): 8951–61.

Yang, Jun, Sunny Sharma, Peter Watzinger, Johannes David Hartmann, Peter Kötter, and Karl-Dieter Entian. 2016. "Mapping of Complete Set of Ribose and Base Modifications of Yeast rRNA by RP-HPLC and Mung Bean Nuclease Assay." Edited by Katrin Karbstein. *PloS One* 11 (12): e0168873.
